# CTLA-4 inhibitors drive colitis through metabolic reprogramming-mediated Treg/Th17 imbalance

**DOI:** 10.64898/2026.01.20.700699

**Authors:** Zhiying Li, Shanna Wu, Rongchen Liu, Runqiu Chen, Fanglin Li, Rong Zhang, Yiting Wang, Chen Chen, Xinhang Zheng, Fengru Qiu, Liying Chen, Yaqi Zhao, Fengyu Du, Likun Gong, Yiru Long

## Abstract

Immune checkpoint inhibitors (ICIs), especially CTLA-4 inhibitors (αCTLA-4), exhibit a high incidence of colitis as an immune-related adverse event (irAE) during cancer treatment, severely limiting patient benefit. Clinically, both treatment interruption and existing intervention drugs for ICI-mediated colitis may compromise antitumor efficacy. However, there is inadequate research on the pathogenesis of ICI-mediated colitis, with findings often conflicting. Here, we first established multiple clinically relevant animal models, including an immuno-humanized ICI-mediated colitis model. Through time-series transcriptomics, we discovered that αCTLA-4-induced colonic toxicity exhibits characteristics ranging from early metabolic reprogramming represented by glycolysis to later immune disorders represented by Th17 responses. By targeting colonic CTLA-4^+^ T cells, αCTLA-4 blocked CD80/CD86-CTLA-4 interaction, thereby activating the PI3K-AKT-mTOR pathway. Subsequently, mTOR mediated metabolic reprogramming in T cells, shifting them from Treg-biased oxidative phosphorylation to Th17-biased glycolysis. The colonic toxicity of αCTLA-4 has also been demonstrated to depend on the PI3K-AKT-mTOR pathway, glycolysis, and Th17 responses. Notably, metformin significantly relieved ICI-mediated colitis by inhibiting mTOR without impeding antitumor efficacy. Collectively, these findings highlighted the metabolic-immune axis in the colonic toxicity of ICI and provided a clinically superior intervention strategy.

## Introduction

Immune checkpoint inhibitors (ICIs), as representative cancer immunotherapies, can restore antitumor immune responses by releasing immune cells from negative regulatory signals. Currently, four targets for ICI therapy—CTLA-4, PD-1, PD-L1, and LAG-3—have been approved, achieving breakthrough progress in the treatment of multiple cancers, such as melanoma and colorectal cancer. Although ICIs typically demonstrate better clinical tolerability relative to conventional treatments such as chemotherapy, they may nonetheless provoke immune-related adverse events (irAE) impacting various organs ^1,2^. Specifically, gastrointestinal toxicity of ICIs constitutes a prevalent and potentially life-threatening irAE, primarily manifesting as colitis characterized by symptoms such as diarrhea, bloody stools, and colonic ulceration. Epidemiological studies indicate that CTLA-4 inhibitors (αCTLA-4) are more likely to induce colitis, with an incidence rate reaching 27% ^1^. When αCTLA-4 is combined with PD-1/PD-L1 inhibitors, the incidence of colitis can rise to 54%, with significantly exacerbated symptoms ^1^. Severe cases may necessitate discontinuation of treatment and adoption of intervention, potentially leading to lifelong exclusion from ICI therapy.

After discontinuing the medication, the patient should initially be treated with the glucocorticoid prednisone. If no improvement is observed, consider initiating therapy with the TNF-α neutralizing antibody infliximab or the integrin α4β7 inhibitor vedolizumab. However, several studies have raised concerns about the efficacy of these three clinical intervention drugs. For instance, researchers found that all three classes of drugs significantly impair the in vivo antitumor activity of ICIs ^3^. Some studies even suggest that integrin α4β7 inhibitors fail to improve ICI-mediated colitis ^4,5^. Clinical retrospective studies have also revealed that these interventions may promote tumor progression ^6,7^. Therefore, the occurrence of ICI-mediated colitis not only leads to discontinuation of treatment, thereby affecting patients’ sustained benefits, but may also compromise cancer patients’ survival due to the use of intervention drugs.

The development of more effective therapeutic interventions requires a deeper understanding of the pathogenesis of ICI-mediated colitis. However, current research into the mechanisms underlying ICI-mediated colitis remains in its infancy, and findings across studies are often inconsistent. Regarding key cells, *Science* reported that Fc-mediated deletion of intestinal regulatory T cells (Tregs) by CTLA-4 inhibitors disrupts intestinal tolerance, thereby triggering colitis ^8^. In contrast, two other studies failed to observe a depletion of intestinal Tregs in mouse models or patients; indeed, an elevation was even noted ^9,10^. Regarding pathogenic factors, researchers proposed that neutralizing tumor necrosis factor-α (TNF-α), interleukin-6 (IL-6) and IL-23 may alleviate colonic toxicity of ICI, though conflicting evidence exists concerning whether such neutralization impacts the antitumour efficacy of ICIs ^4,10,11^. In other respects, several studies have suggested that the gut microbiota influences the development of ICI-mediated colitis ^8,12,13^, though the precise mechanisms remain unclear.

Therefore, in this study, we focused on CTLA-4 inhibitors with high frequency of colitis to research the mechanisms responsible for ICI-induced colitis, and then investigated novel intervention strategies for balancing the effectiveness and toxicity of ICIs.

## Results

### Establishment of DSS-based mouse models of ICI-mediated colitis

Laboratory specific pathogen free (SPF) mice generally possess tolerance to ICI therapy, making it challenging to induce systemic toxicity with directly administered various ICI drugs. We first tested the αCTLA-4 9H10 and IgG isotype control in C57BL/6 mice bearing B16F10 melanoma tumors. After continuous administration, we assessed diarrhea/bloody stool scores (DAI), colon length, and colonic pathology. The results showed that the CTLA-4 inhibitor did not cause any observable colonic injury (Fig.S1).

The above results indicated that basic induction conditions are required when establishing an ICI-mediated colitis mouse model. DSS is a chemical commonly used to induce a colitis mouse model. By providing mice with DSS drinking, it pre-damages the intestinal barrier, thereby helping to overcome the mice’s tolerance to ICIs. We first screened the DSS drinking concentrations, including 2.5%, 3%, 4%, and 5%, in B16F10-bearing mice (Fig.S2). Following intraperitoneal injection of the αCTLA-4 9H10 and IgG, we compared differences between the two groups in aspects of body weight, DAI score, and survival (Fig.S2A-B). We found that αCTLA-4 resulted in significant weight loss in mice compared to IgG in the 2.5% and 4% groups. Similarly, administration of αCTLA-4 in the 2.5% and 4% groups resulted in markedly elevated DAI scores. Regarding mouse survival, results showed that 3%, 4%, and 5% DSS drinking all resulted in mouse mortality, except for the 2.5% concentration (Fig.S2C). In the 4% group, the CTLA-4 inhibitor caused 50% mortality. Therefore, considering the model window size, as well as animal welfare and model stability, we selected αCTLA-4 administering with 2.5% DSS drinking to cause ICI-mediated colitis in mice.

Besides, based on this DSS condition, we also established mouse models of αCLTA-4/αPD-1 combination-mediated colitis and PD-1 inhibitor-mediated colitis (Fig.S3). Two monotherapies and the αCLTA-4/αPD-1 combination caused significant weight loss and increased DAI scores (Fig.S3A-B). Furthermore, the αCLTA-4/αPD-1 combination group exhibited a further trend toward reduced colon length compared to the two monotherapy groups (Fig.S3C), consistent with clinical manifestations.

### Establishment of immuno-humanized mouse models of ICI-mediated colitis

To further mimic the development of ICI-mediated colitis in a human-derived system, we attempted to reconstruct the human immune system by reinfusing human peripheral blood mononuclear cells (hPBMCs) into severely immunodeficient NCG mice. On this basis, the mice were administered the marketed αCLTA-4 ipilimumab and αPD-1 nivolumab.

In previous studies, researchers initiated ICIs treatment concurrently with hPBMCs reinfusion. However, we found no differences between the PBMC+IgG and PBMC+ICI groups in body weight, DAI, or colon length, indicating that this approach fails to induce colitis (Fig.S4A). Additionally, the concurrent tumor implantation and hPBMC reinfusion regimen also failed (Fig.S4B). Given that the classic immune reconstitution cycle for hPBMCs takes approximately 21 days, we hypothesized that administering ICIs after completing immune reconstitution is critical for model success. In this strategy, compared to the IgG group, administration of αCTLA-4/αPD-1 resulted in significant weight loss in mice, markedly elevated DAI scores, and a noticeable reduction in colon length (Fig.S4C). These findings indicated the successful establishment of an immuno-humanized ICI-mediated colitis model, which could be utilized for subsequent research.

### Colonic toxicity phenotype induced by CTLA-4 inhibitors

Upon the DSS-based mouse model of αCTLA-4-induced colitis, we further systematically analyzed the colonic toxicity of CTLA-4 inhibitors. Regarding body weight, the αCTLA-4 group exhibited a decline starting on Day 4, reached its nadir by Day 7, and then saw a progressive recovery, in contrast to the IgG group (Fig.1A-B). CTLA-4 inhibitors caused an approximately 1 cm shortening of the colon, suggesting longitudinal contraction due to thickening of the colonic muscularis (Fig.1C). Hematoxylin and eosin staining (H&E) results revealed extensive crypt loss/atrophy in the colon following αCTLA-4 treatment, accompanied by marked inflammatory cell infiltration and significant submucosal edema (Fig.1D).

**Fig. 1.**
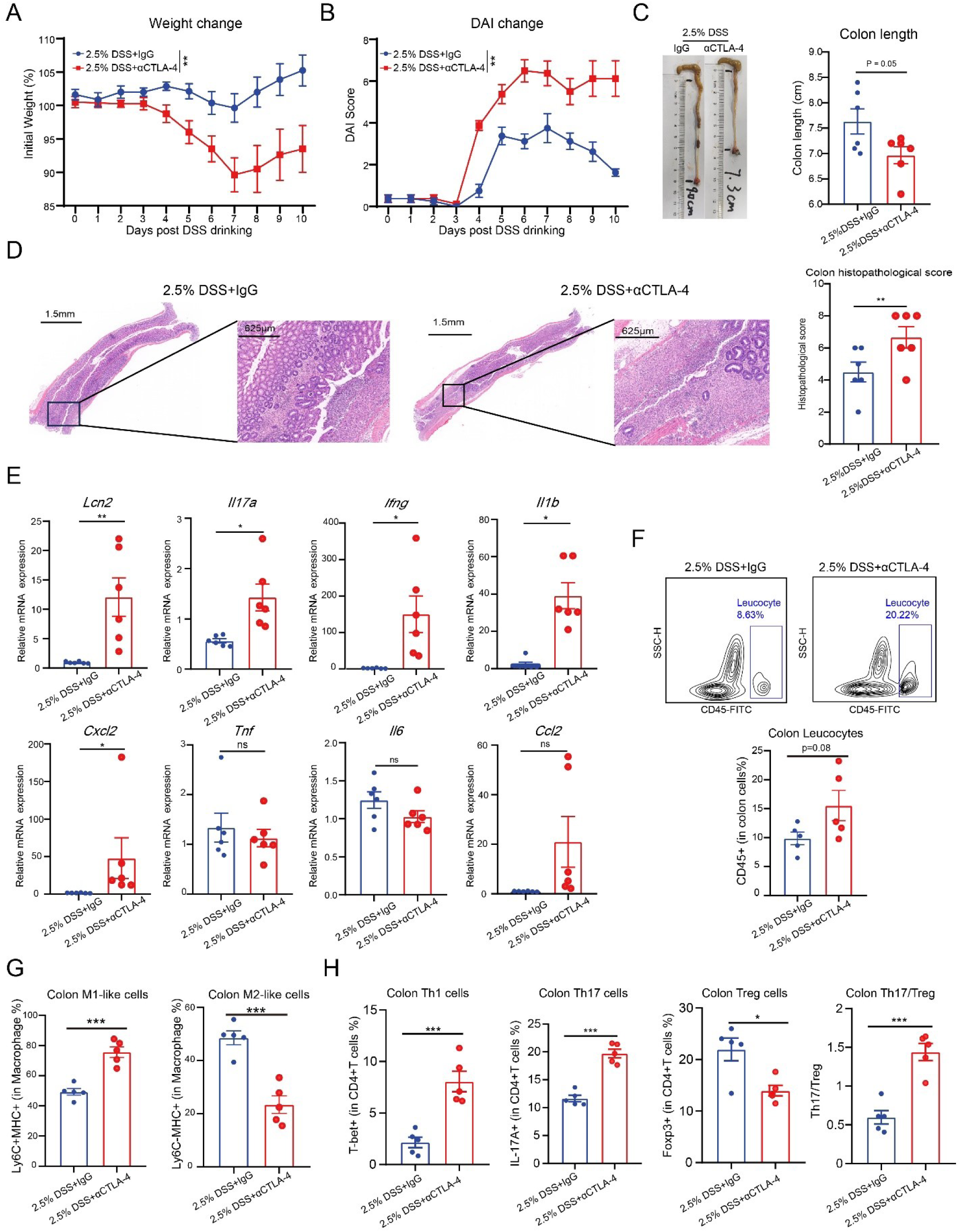
Administration of αCTLA-4 disordered the colonic immune microenvironment. B16F10 tumor-bearing C57BL/6 mice were provided with 2.5% DSS drinking water from D0 to D3. On D0, D3, D6, and D9, mice received intraperitoneal injections of 200 μg/mouse αCTLA-4 9H10 or isotype control IgG. (**A-B**) Body weight monitoring and DAI score monitoring were performed (n=6). (**C**) Representative colon photographs and colon length statistics (n=6). (**D**) Representative colon H&E staining results and statistical analysis of colonic pathology scores (n=6). (**E**) Transcription levels of multiple inflammatory and chemokines in mouse colon tissues were detected (n=6). (**F**) Representative gating and statistical analysis of leukocytes in the colon tissues (n=5). (**G**) Flow cytometric analysis of M1-like and M2-like macrophages in the colon tissues (n=5). (**H**) Flow cytometric analysis of CD4+ T cell subpopulations in the colon tissues (n=5). The data are presented as the mean ± SEM. * p < 0.05; ** p < 0.01; *** p < 0.001; ns not significant by ns not significant by unpaired t test.

Subsequently, we systematically analyzed alterations in the colonic immune microenvironment. CTLA-4 inhibitors resulted in approximately tenfold upregulation of colonic lipocalin 2 (LCN-2) levels, a biomarker for clinical ulcerative colitis (Fig.1E). Colonic IFN-γ, IL-17A, and IL-1β levels were significantly upregulated, but no significant changes were observed in IL-6, TNF-α, and other markers (Fig.1E). Multicolor flow cytometry analysis of colonic lamina propria immune cells revealed that administration of CTLA-4 inhibitors increased white blood cell infiltration in the colon by approximately 1.5-fold (Fig.1F). Further, the proportion of pro-inflammatory M1-like macrophages significantly increased, while the proportion of anti-inflammatory M2-like macrophages significantly decreased (Fig.1G). The percentage of Th1/Th17 cells in the colon significantly increased, according to the analysis of CD4+ T cell subsets (Fig.1H). Moreover, Th17 cells have emerged as the most prevalent CD4+ T cell subgroup in the colon, accounting for up to 20% of the total. These findings are also consistent with the upregulation of IL-17A and IFN-γ detected at the mRNA level. Further, the Th17/Treg ratio significantly increased following administration of αCTLA-4, suggesting disruption of the colonic immune homeostasis, shifting it toward a Th17-biased inflammatory environment.

Given the controversy in previous studies regarding whether CTLA-4 inhibitors influence colitis development via Tregs, we selected the non-Treg-deleting 9H10 antibody and the Treg-deleting 9D9 antibody for modeling (Fig.S5). Based on body weight changes and DAI scores, no significant differences were observed. Therefore, we speculate that the colitis toxicity of CTLA-4 inhibitors may correlate with CTLA-4 signaling, rather than the antibody-mediated Treg deletion.

### Time-series transcriptome sequencing analysis of αCTLA-4-mediated colitis

To investigate the pathogenesis of αCTLA-4-mediated colitis, we dissected colon tissues from mice at Days 4, 7, and 11 based on the aforementioned model (Fig.2A). RNA sequencing (RNA-seq) was performed to analyze gene expression alterations during the early, intermediate, and late phases of αCTLA-4 administration. In the early stages, KEGG enrichment analysis revealed that, compared to the IgG control, αCTLA-4 unexpectedly induced alterations primarily in metabolic pathways—including glycolysis, fatty acid metabolism, and amino acid metabolism—while showing no obvious changes in immune-related pathways (Fig.2B). In later stages, αCTLA-4 primarily induced alterations in immune-related pathways, such as the IL-17 pathway, Th17 differentiation, PI3K-AKT pathway, HIF-1 signaling pathway, and inflammatory bowel disease pathway (Fig.2B).

**Fig. 2.**
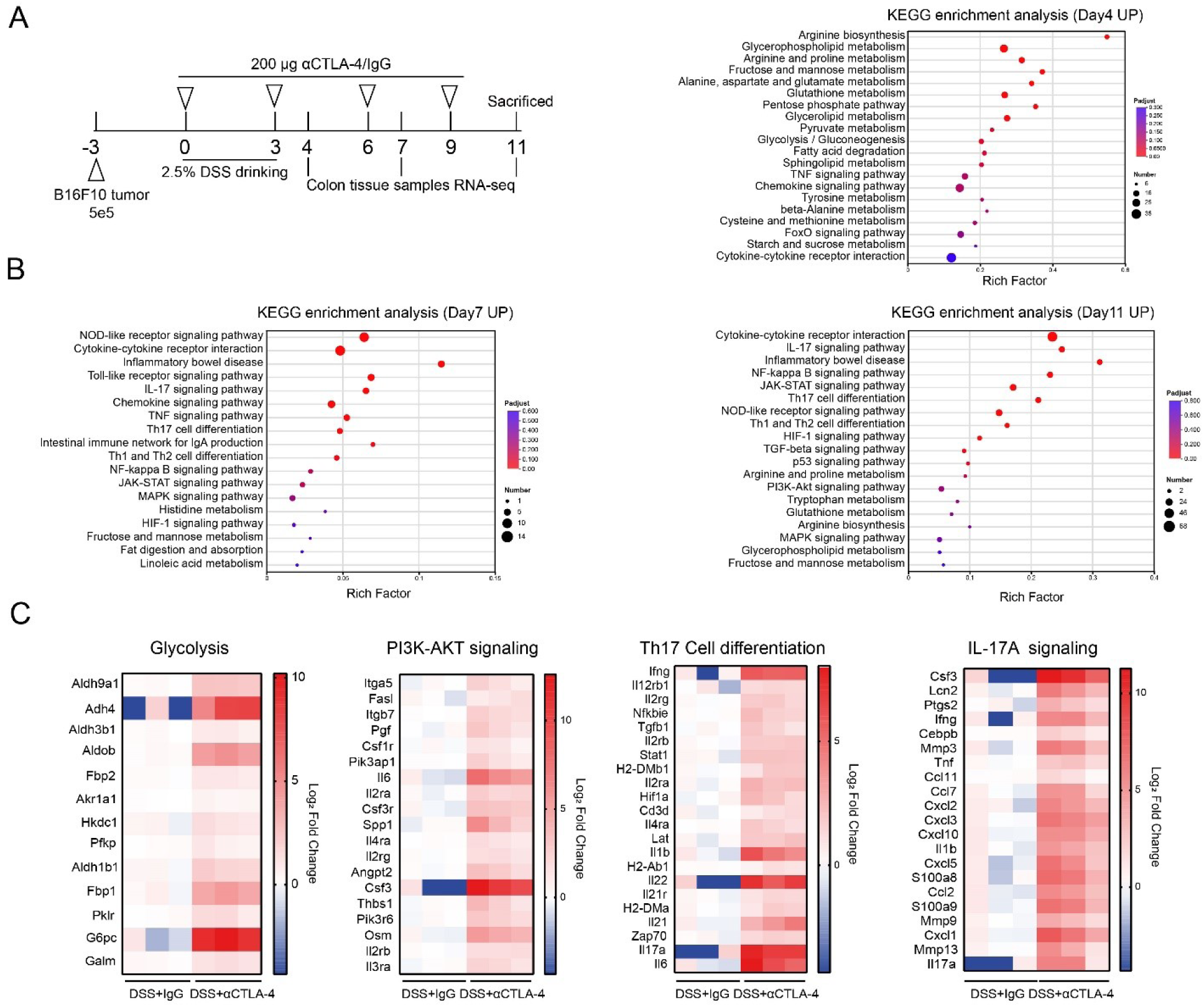
Administration of αCTLA-4 was accompanied by a shift from colonic metabolic alterations to immune alterations. (**A**) Early/middle/late point sampling roadmap. (**B**) KEGG pathway enrichment analysis of differentially expressed genes in the DSS+αCTLA-4 group at three time points compared to the DSS+IgG group. (**C**) Heatmap of gene expression levels related to glycolysis, PI3K-AKT pathway, Th17 cell differentiation, and IL-17A signaling in the DSS+IgG group and DSS+αCTLA-4 group.

In previous studies, researchers observed that the endogenous ligands CD80/CD86 transmit signals to CTLA-4, thereby inhibiting the activation of the PI3K-AKT-mTOR pathway^14^. In metabolic research, mTOR is a well-known key regulatory molecule whose activation promotes pathways such as glycolysis and fatty acid synthesis. In recent years, researchers have discovered that glycolysis could promote Th17 cell differentiation and function, while Treg cells prefer oxidative phosphorylation^15,16^. We then analyzed relevant pathways in the RNA-seq data and found that αCTLA-4 consistently upregulated genes associated with glycolysis, the PI3K-AKT pathway, Th17 differentiation, and IL-17A signaling (Fig.2C).

These findings suggested that αCTLA-4-mediated colitis may arise from early metabolic reprogramming driving alterations in the immune microenvironment.

### αCTLA-4 promoted the colonic PI3K-AKT-mTOR, glycolysis, and IL-17 pathway

To validate the above hypothesis, we first examined the transcriptional expression of associated genes. Results showed that IL-17 pathway-related gene expression was predominantly upregulated in the colon at the late phase of treatment, especially the IL-17A (Fig.3A). At the early phase, expression of key glycolytic enzymes such as fructose-1,6-bisphosphatase 1 (FBP1) was also largely increased (Fig.3B), consistent with the RNA-seq findings.

**Fig. 3.**
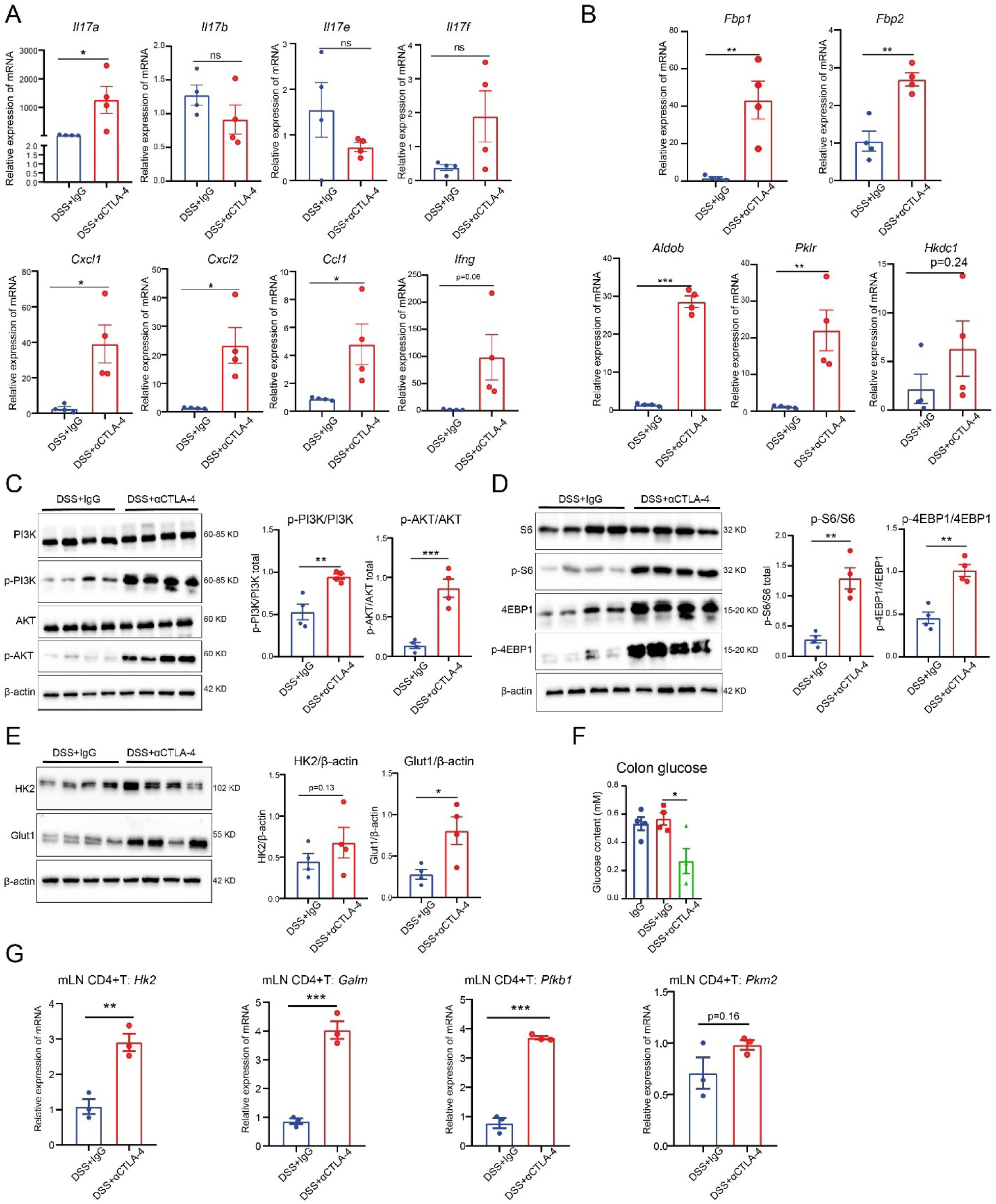
CTLA-4 inhibitor amplified colonic PI3K-AKT-mTOR, glycolysis, and IL-17 signaling pathways. B16F10 tumor-bearing C57BL/6 mice were provided with 2.5% DSS drinking water from D0 to D3. On D0, D3, D6, and D9, mice received intraperitoneal injections of 200 μg/mouse αCTLA-4 9H10 or isotype control IgG. (**A**) Transcriptional levels of IL-17A pathway-related molecules at the late stage in the colon of mice administered CTLA-4 inhibitor (n=4). (**B**) Transcriptional levels of glycolysis-related molecules at the early stage in the colon of mice administered CTLA-4 inhibitor (n=4). (**C**) Western blot analysis of total and phosphorylated PI3K and AKT proteins in mouse colon tissues (n=4). (**D**) Western blot analysis of total and phosphorylated S6 and 4EBP1 proteins in mouse colon tissues (n=4). (**E**) Western blot analysis of total HEK2 and Glut1 proteins in mouse colon tissues (n=4). (**F**) Glucose content measurement in mouse colon tissues at the model endpoint (n=4). (**G**) Gene expression of *Hk2*, *Galm*, *Pfkb1*, and *Pkm2* in CD4+ T cells sorted from mouse mLNs at the model endpoint (n=3). The data are presented as the mean ± SEM. * p < 0.05; ** p < 0.01; *** p < 0.001; ns not significant by unpaired t test or one-way ANOVA followed by Tukey’s multiple comparisons test.

At the protein level, we performed Western blot (WB) analysis on mouse colon tissues at the model endpoint and found that following αCTLA-4 administration, the phosphorylation levels of PI3K, AKT, S6, and 4EBP1 were significantly elevated (Fig.3C-D), indicating marked activation of the PI3K-AKT-mTOR pathway. Moreover, the key glycolytic proteins glucose transporter 1 (Glut1) and Hexokinase-2 (HK2) were both upregulated (Fig.3E). Correspondingly, glucose levels in colonic tissues decreased significantly (Fig.3F).

Subsequently, to validate that these pathway alterations occurred in CD4+ T cells, we isolated CD4+ T cells from the mesenteric lymph nodes (mLNs) at the model endpoint. At the transcriptional level, we found that αCTLA-4 indeed led to a significant upregulation of glycolysis-related gene expression in CD4+ T cells (Fig.3G).

The above results further validated that CTLA-4 inhibitors activate the PI3K-AKT-mTOR, glycolysis, and IL-17 pathways in the colon.

### αCTLA-4-mediated colitis was dependent on Th17 cells

To validate that the colonic toxicity of αCTLA-4 depends on Th17 cells, we administered a neutralizing antibody against IL-17A—a key effector of Th17 cells—to mice in the DSS-based αCTLA-4-mediated colitis model. Based on body weight, DAI scores, and colon length metrics, neutralizing IL-17A significantly alleviated colonic toxicity (Fig.4A-B). Moreover, IL-17A neutralization did not impair the antitumor activity of αCTLA-4 (Fig.4C-D). Neutralizing IL-17A also effectively improved αCTLA-4-induced colonic pathological damage (Fig.4E).

**Fig. 4.**
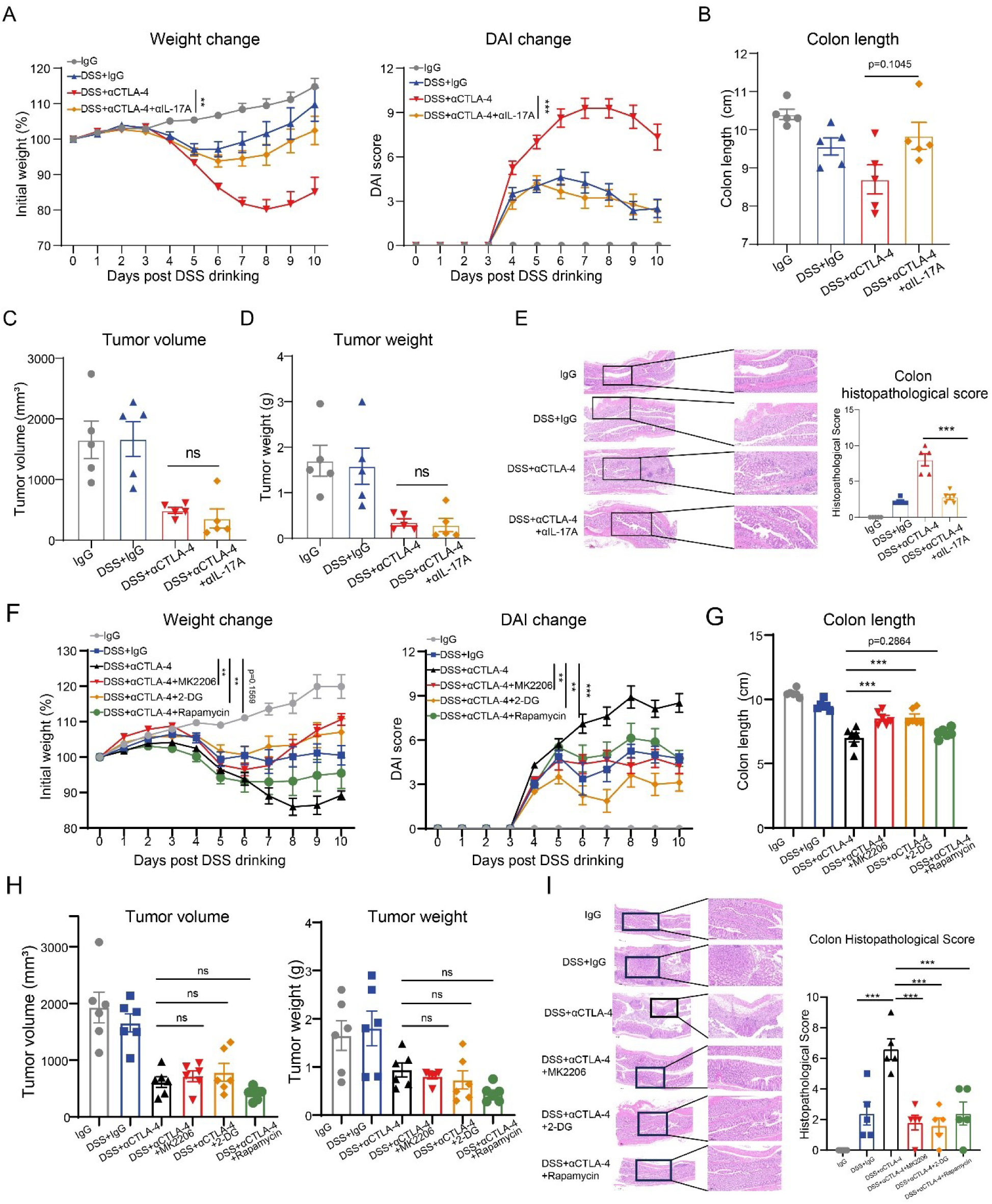
Colonic toxicity of αCTLA-4 depended on PI3K-AKT-mTOR, glycolysis, and IL-17A signaling. (**A-E**) In the DSS-based αCTLA-4-induced colitis model, anti-mouse IL-17A neutralizing antibody (200 μg/mouse) was administered intraperitoneally at D0, D3, D6, and D9 (n=5). Body weight monitoring (A), DAI score monitoring (A), colon length measurement (B), endpoint tumor volume (C) and tumor weight (D) detection, representative colon H&E results and pathological score statistics (E). (**F-I**) In the DSS-based αCTLA-4-induced colitis model, MK2206 (p.o.; 30 mg/kg/day), Rapamycin (i.p.; 5 mg/kg/day), and 2-DG (i.p.; 500 mg/kg/day) were administered from D0 to D10 (n=6). Body weight monitoring (F), DAI score monitoring (F), colon length measurement (G), endpoint tumor volume (H) and tumor weight (H) detection, representative colon H&E results and pathological score statistics (I). The data are presented as the mean ± SEM. ** p < 0.01; *** p < 0.001; ns not significant by one-way ANOVA followed by Tukey’s multiple comparisons test.

These findings suggested that the colonic toxicity of CTLA-4 inhibitors was indeed dependent on CD4+ T cells, particularly Th17 cells.

### Colonic toxicity of αCTLA-4 depended on the PI3K-AKT-mTOR pathway and glycolysis

To validate that the colonic toxicity of αCTLA-4 depended on the PI3K-AKT-mTOR pathway and glycolysis, we administered the AKT inhibitor MK2206, the mTOR inhibitor rapamycin, and the glycolysis inhibitor 2-DG to mice in the DSS-based αCTLA-4-mediated colitis model. Results showed that all three inhibitors significantly improved weight loss, DAI score elevation, and colon length shortening induced by αCTLA-4 (Fig.4F-G). And they did not affect the anti-tumor efficacy of αCTLA-4 (Fig.4H). In regard to colonic pathology, all three inhibitors significantly alleviated the colonic damage (Fig.4I).

Furthermore, inhibition of AKT/mTOR and glycolysis both significantly reduced colonic IL-17A expression, while AKT/mTOR inhibition could downregulate colonic glycolysis-related gene expression.

Based on the above findings, it was primarily validated that αCTLA-4 amplified Th17-mediated colonic inflammation by regulating PI3K-AKT-mTOR-mediated CD4+ T cell metabolic reprogramming.

### Evaluation of the effect of metformin on ICI-mediated colitis

Based on the above hypothesis, we aimed to select a clinically available drug that targets the PI3K/AKT/mTOR pathway and glycolysis, to evaluate its efficacy in improving αCTLA-4-mediated colitis. This approach sought to ensure the drug’s rapid availability for clinical patients. Metformin, a clinically used diabetes drug, can activate AMPK to inhibit the PI3K/AKT/mTOR pathway, thereby suppressing glycolysis. Given its generally positive safety profile and low cost, we plan to conduct a systematic evaluation of its effects on the colonic toxicity and antitumor efficacy of αCTLA-4.

First, during administration of αCTLA-4, mice were treated daily via oral gavage with either 100 mg/kg or 200 mg/kg of metformin hydrochloride solution for the intervention study. Metformin nearly completely eliminated the weight loss and DAI scores increasing induced by αCTLA-4 (Fig.5A). Moreover, even at the relatively low dose of 100 mg/kg, metformin demonstrated potent ameliorative effects. Metformin also exhibited a dose-dependent reversal of colonic shortening (Fig.5B). Based on the clinical marker LCN-2 level, a 200 mg/kg dose of metformin nearly completely resolved αCTLA-4-mediated colitis, demonstrating superior efficacy (Fig.5C). This finding was further validated at the histopathological level, where treated mice exhibited colonic tissue structures more closely resembling normal colonic architecture (Fig.5D).

**Fig. 5.**
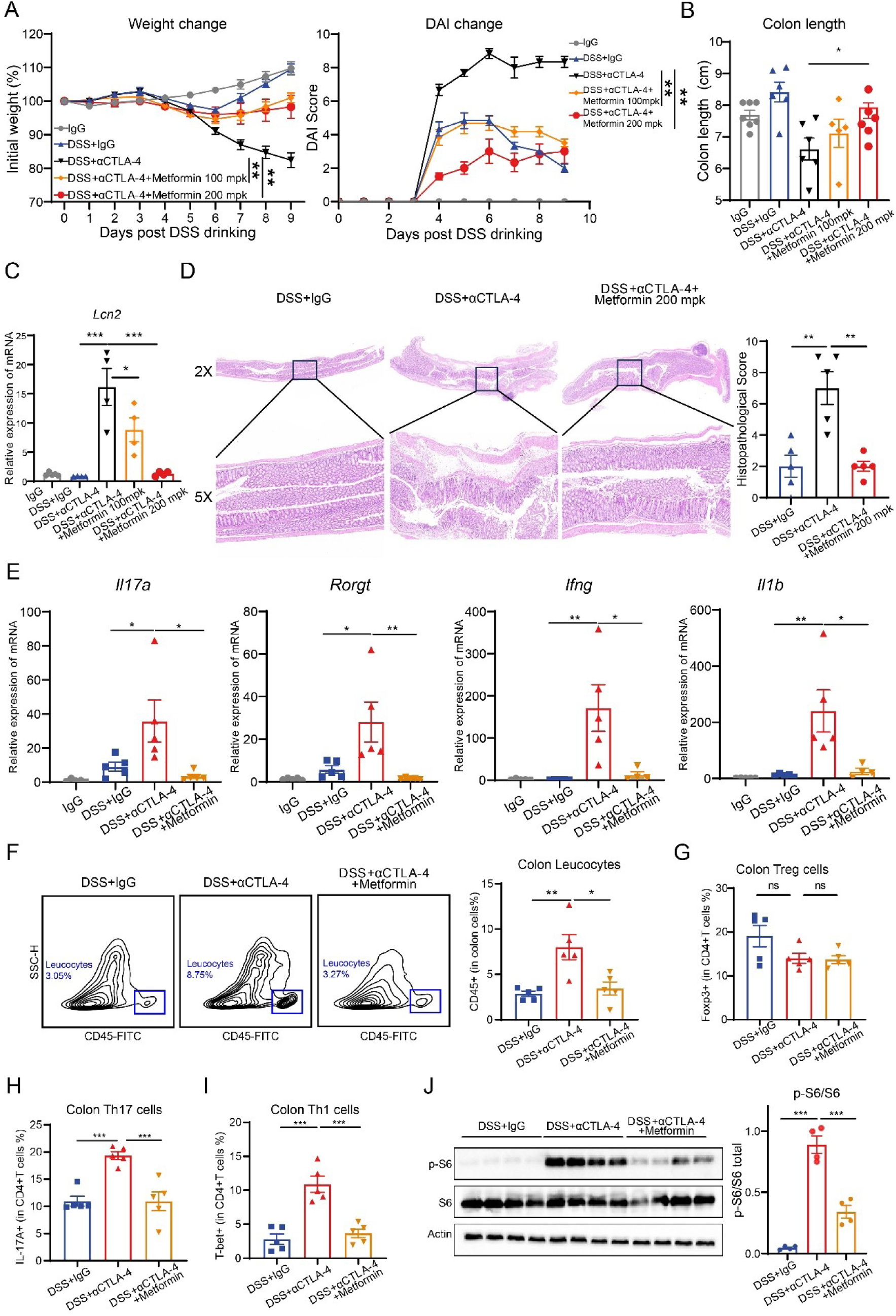
Metformin could effectively alleviate CTLA-4 inhibitor-mediated colitis. In the DSS-based αCTLA-4-induced colitis model, metformin (p.o.; 100 or 200 mg/kg/day) was administered from D0 to D10 (n=5-7). (**A**) Body weight monitoring and DAI score monitoring were performed (n=5-7). (**B**) Colon length statistics (n=5-7). (**C**) Transcriptional levels of *Lcn2* (n=4). (**D**) Representative colon H&E results and pathological score statistics (n=5). (**E**) Detection of colonic *Il17a*, *Rorgt*, *Ifng*, and *Il1b* transcription levels (n=5). (**F**) Representative gating and statistical analysis of leukocytes in the colon tissues (n=5). (**G-I**) Flow cytometric analysis of CD4+ T cell subpopulations in the colon tissues (n=5). (**J**) Western blot analysis of total and phosphorylated S6 proteins in mouse colon tissues (n=4). The data are presented as the mean ± SEM. * p < 0.05; ** p < 0.01; *** p < 0.001; ns not significant by one-way ANOVA followed by Tukey’s multiple comparisons test.

Subsequently, we tested whether metformin improved the indicators in the hypothesis. At the transcriptional level, metformin administration nearly completely downregulated the expression of αCTLA-4-mediated Th17-related markers IL-17A and RORγT, while also suppressing levels of inflammatory mediators such as IFN-γ and IL-1β (Fig.5E). In the colonic microenvironment, metformin administration significantly suppressed leukocyte infiltration (Fig.5F). It had no apparent effect on Treg proportion but inhibited the colonic infiltration of Th17 and Th1 cells (Fig.5G-I). At the protein level, metformin administration significantly downregulated S6 phosphorylation (Fig.5J), thereby inhibiting the activation of the colonic mTOR pathway.

Given that, in addition to CTLA-4, the PD-1/PD-L1 pathway has also been reported in recent years to participate in regulating glycolysis, we hypothesized that metformin may be broadly applicable for improving ICI-mediated colitis. In the aforementioned mouse models of αCLTA-4/αPD-1 combination-mediated colitis and PD-1 inhibitor-mediated colitis, metformin also significantly eliminated the weight loss, DAI scores increasing and colonic shortening (Fig.S6A-C). Similarly, metformin did not affect the antitumor efficacy of the αCLTA-4/αPD-1 combination or PD-1 inhibitor monotherapy (Fig.S6D).

Taken together, metformin could inhibit the mTOR pathway and Th17 responses in the colon, demonstrating potent improvement in ICI-mediated colitis.

### Comparison of the effects of metformin and the clinical intervention drug meprednisone

To further evaluate the impact of metformin on the antitumor efficacy of CTLA-4 inhibitors and to conduct a head-to-head comparison with the clinically applied glucocorticoid drug meprednisone (also called methylprednisolone), we performed experiments in both non-colitis tumor-bearing models and colitis tumor-bearing models. In non-colitis tumor models, metformin administration did not affect the efficacy of CTLA-4 inhibitors as measured by endpoint tumor volume and weight, whereas meprednisone nearly completely eliminated the efficacy (Fig.6A).

**Fig. 6.**
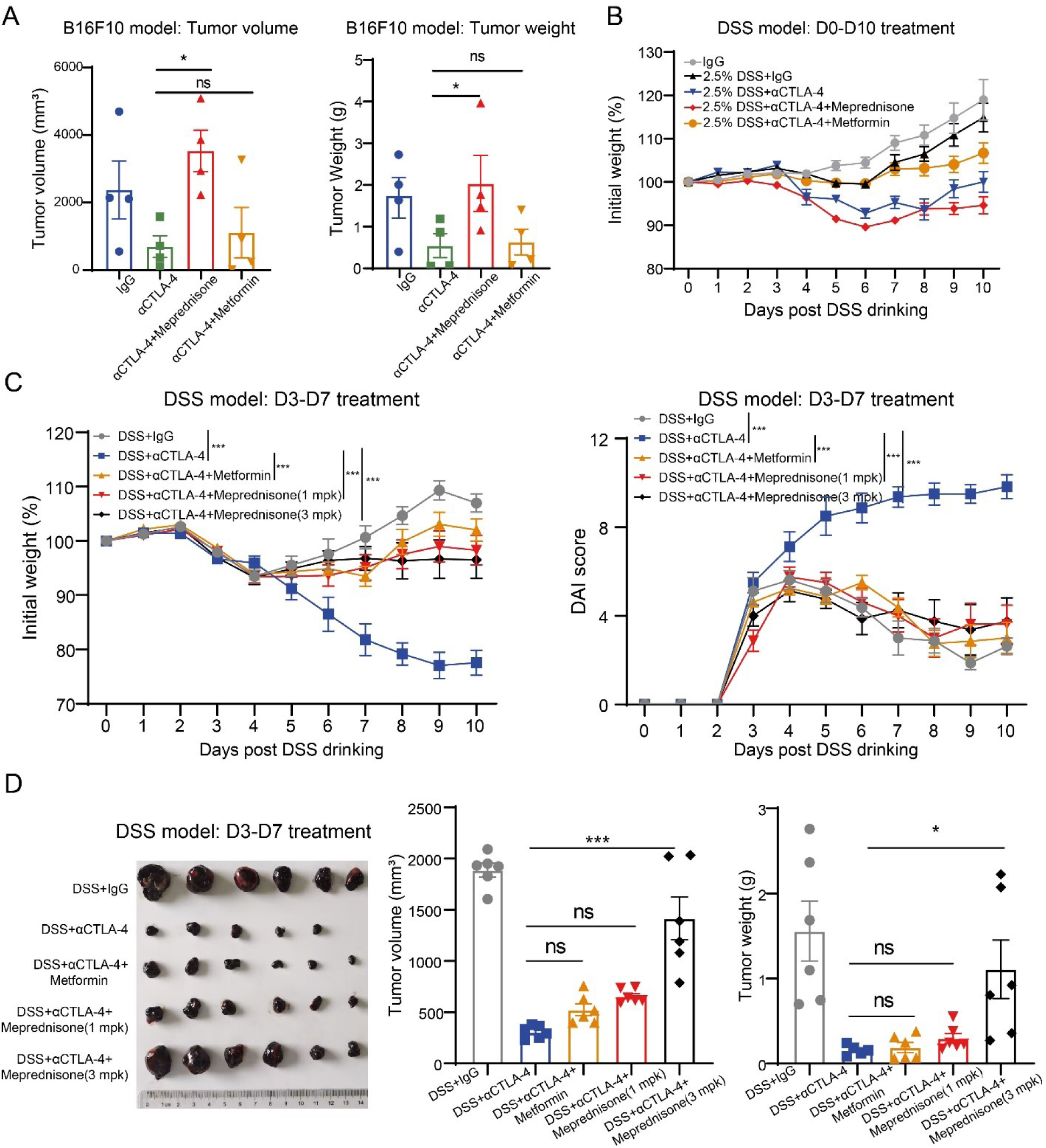
Comparison of metformin and meprednisone in improving αCTLA-4-mediated colitis. (**A**) In the non-colitis tumor-bearing model, mice were administered 1 mg/kg meprednisone and 200 mg/kg metformin from days 3 to 10. Tumor volume and weight were measured at sacrifice on day 11 (n=4). (**B**) In the colitis tumor-bearing model, mice received 1 mg/kg meprednisone and 200 mg/kg metformin from days 0 to 10, with body weight changes monitored (n=6). (**C-D**) In the colitis tumor-bearing model, mice were administered 1 mg/kg or 3 mg/kg meprednisone and 200 mg/kg metformin from days 3 to 7. Changes in body weight and DAI scores were monitored, and tumor volume and weight were measured at the endpoint (n=6). The data are presented as the mean ± SEM. * p < 0.05; *** p < 0.001; ns not significant by one-way ANOVA followed by Tukey’s multiple comparisons test.

Subsequently, in the ICI-induced colitis model, we observed that when administered from D0 to D10, metformin effectively mitigated weight loss (Fig.6B). However, mice demonstrated intolerance to meprednisone, which even exacerbated weight loss (Fig.6B). To effectively alleviate αCTLA-4-mediated colitis while comparing metformin and meprednisone, we adjusted the dosing regimen to administer the drugs from D3 to D7 (Fig.6C). Results showed that mice tolerated meprednisone well, and it effectively mitigated the weight loss and DAI scores increasing induced by CTLA-4 inhibitors (Fig.6C). The ameliorative effect of metformin was comparable to that of meprednisone. Notably, analysis of tumor volume and weight revealed that meprednisone dose-dependently interfered with the antitumor efficacy of αCTLA-4, whereas metformin did not affect the efficacy (Fig.6D).

Taken together, unlike the clinical drug meprednisone, metformin could significantly ameliorate the colonic toxicity of αCTLA-4 without compromising their antitumor efficacy. Metformin may help ICI drugs achieve a balance between efficacy and toxicity, demonstrating great potential for clinical application.

### Single-cell transcriptome sequencing analysis of patient samples with ICI-mediated colitis

To further validate whether our hypothesis extends to humans, we collected open-source single-cell RNA sequencing (scRNA-seq) data from the GEO database (GSE144469). This dataset includes colon tissue samples from 8 healthy volunteers, 8 melanoma patients treated with ICIs who developed colitis, and 6 melanoma patients treated with ICIs who did not develop colitis. CD4+ T cell subsets were isolated and further subdivided into nine populations, including Treg, Tnaive, Tfh, Th17, Th1, Tcytotoxic, Tcycling, and others (Fig.7A). Analysis of CTLA-4 gene expression levels across various subsets similarly revealed its highest expression in colonic Treg cells, with detectable expression also observed in other T cell types such as Th17 cells (Fig.7B-C).

**Fig. 7.**
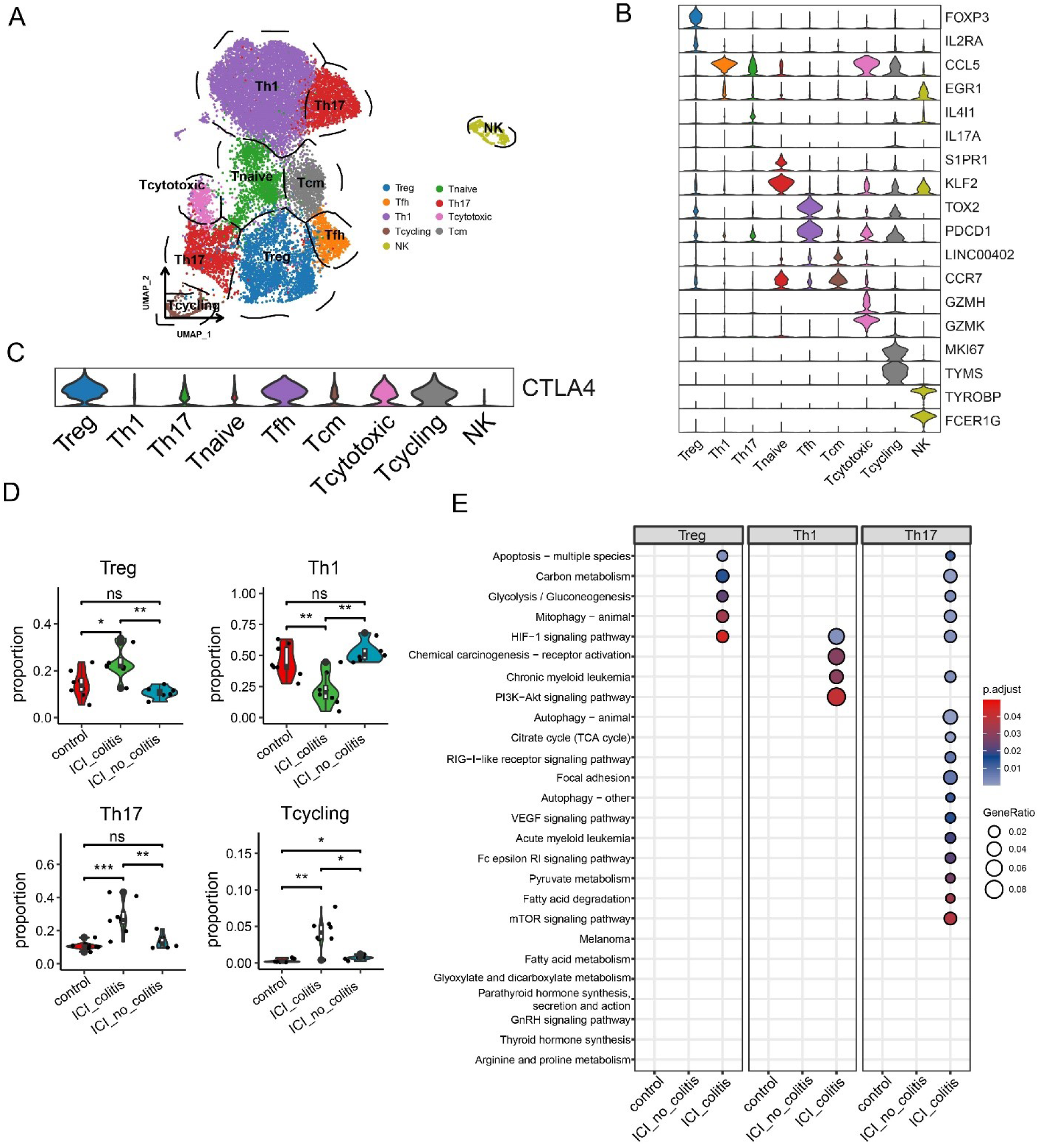
scRNA-seq analysis of patient samples with ICI-mediated colitis. Re-analysis of CD4+ T-related cells in clinical healthy control samples, ICI_colitis samples, and ICI_no_colitis samples (n=6-8). (**A**) Annotation of CD4+ T-related cells. (**B**) Gene expression levels characteristic of each cell subpopulation. (**C**) CTLA-4 gene expression levels in various cell subsets.

Subsequently, we analyzed the proportions of various CD4+ T cell subsets across different groups. Results revealed a significant increase in Th17 cells among patients with ICI-mediated colitis, accompanied by an increase in Tregs, consistent with our previous findings (Fig.7D).

Furthermore, no increase in Th1 cells was observed in patients with ICI-mediated colitis, suggesting that Th17 rather than Th1 cells should remain the primary focus among pro-inflammatory T cells (Fig.7D). KEGG enrichment analysis revealed that the glycolysis was highly enriched in Treg and Th17 cells from ICI-mediated colitis patients (Fig.7E). The PI3K-AKT pathway, mTOR signaling pathway, and the associated HIF-1 pathway were also highly expressed in Th17 cells from patients with ICI-mediated colitis (Fig.7E).

These findings were consistent with our experimental results, suggesting that our hypothesis may also hold true in real clinical patients and the clinical application potential for metformin.

## DISCUSSION

Our research has revealed that metabolism-driven Treg/Th17 imbalance plays a crucial role in the development of colitis mediated by CTLA-4 inhibitors, with metformin emerging as an ideal candidate intervention drug.

When establishing the mouse model of αCTLA-4-mediated colitis, we found that SPF mice exhibited good tolerance to CTLA-4 inhibitors, with direct administration not causing colonic injury, which is consistent with previous reports^8^. In our study, disruption of the intestinal barrier via DSS drinking or GVHD induced by hPBMCs reinfusion helped overcome mouse tolerance to ICIs. This also suggests that certain predisposing factors exist for the occurrence of colitis adverse reactions in clinical cancer patients receiving ICIs treatment. We hypothesize that the patient’s genetic background, prior medication history, gut health status, living environment, history of microbial infections, and other factors may all be predisposing factors for ICI-mediated colitis.

Metformin, as a clinical first-line oral hypoglycemic agent, has demonstrated anti-inflammatory and metabolic reprogramming activities in various diseases beyond its classic blood glucose-regulating effects, such as cancers, metabolic syndrome, and autoimmune diseases by inhibiting the mTOR pathway through AMPK activation^17,18^. Although metformin has not yet been observed to be used for treating ICI-mediated colitis irAE, studies have reported that metformin can be used to treat ulcerative colitis. For example, researchers found that metformin can suppress excessive activation of the STAT3 by downregulating acetyl-CoA levels, thereby alleviating colitis ^19^. Besides, metformin has also been reported to improve colitis by regulating the gut microbiota ^20^. In our study, metformin could suppress ICI-induced gut metabolic-immune dysregulation by inhibiting the mTOR pathway, thereby restoring the gut Treg/Th17 balance.

It is noteworthy that although clinical guidelines recommend glucocorticoids, TNF-α inhibitors, and integrin α4β7 inhibitors for the management of ICI-mediated colitis, all three drugs have been reported to impair the antitumor efficacy of ICIs. This is also consistent with our findings that meprednisone affects the efficacy of CTLA-4 inhibitors in multiple models. Therefore, we consider metformin an ideal intervention for ICI-mediated colitis, capable of balancing the colonic toxicity of ICIs with their antitumor efficacy. Considering that despite metformin’s favorable clinical safety profile, certain side effects do exist in practice. Simultaneously, to more precisely address the “on-target off-tumor” toxicity of CTLA-4 inhibitors, we envisioned creating a novel antibody drug conjugates (ADC), the CTLA-4 antibody-metformin conjugate, by linking metformin to the CTLA-4 antibody. This approach will enable metformin to be precisely delivered alongside the CTLA-4 inhibitor to the sites where side effects occur.

To further explore the potential of metformin in preventing or treating clinical ICI-associated colitis, considering its broad clinical use for diabetes, we plan to conduct a retrospective study. This study will collect information on three groups of patients undergoing ICI therapy: 1. Patients with underlying diabetes who have been on long-term metformin therapy prior to ICI treatment. 2. Patients with underlying diabetes who have discontinued metformin within the past year before starting ICI therapy. 3. Patients without diabetes before starting ICI therapy. We will compare and analyze the incidence of colitis and other irAEs among these three groups.

In summary, based on the establishment of multiple clinically relevant ICI-mediated colitis mouse models, this study reveals that the colonic toxicity of CTLA-4 inhibitors primarily occurs through metabolic reprogramming-mediated Treg/Th17 imbalance driven by the PI3K-AKT-mTOR pathway. Metformin can improve ICI-associated colitis by restoring colonic metabolic-immune homeostasis through mTOR inhibition, without affecting antitumor efficacy. The metabolism-immunity axis represents a novel mechanism underlying ICI-related irAEs, and metformin holds immense clinical potential as an ideal intervention drug.

## METHODS

### Mice

Male six– to eight-week-old C57BL/6 mice were purchased from Vital River. NCG mice were purchased from GemPharmatech Co., Ltd. All mice were maintained under specific pathogen-free (SPF) conditions in the animal facility of the Shanghai Institute of Materia Medica, Chinese Academy of Sciences. Animal care and experiments were performed in accordance with the Shanghai Institute of Materia Medica, Chinese Academy of Sciences, using protocols approved by the Institutional Laboratory Animal Care and Use Committee (IACUC).

### Cells

B16F10 and RKO cell lines were purchased from Procell. B16-F10 were cultured in Dulbecco’s Modified Eagle Medium (DMEM; Meilunbio) supplemented with 1% penicillin/streptomycin (P/S; Invitrogen) and 10% heat-inactivated fetal bovine serum (FBS; Life Technologies). RKO cells were cultured in Minimum Essential Medium (MEM; Meilunbio) supplemented with 1% P/S and 10% FBS. Human PBMCs were purchased from Milestone. All cells were cultured at 37°C in a 5% CO_2_ humidified atmosphere.

### Establishment of a DSS-based αCTLA-4-mediated colitis

At Day –3, the mice were randomly grouped and injected subcutaneously with 5×10^5^ of B16F10 tumor cells in the right forelimb. B16F10 tumor-bearing C57BL/6 mice were provided with DSS (60316ES76, Yeasen) in drinking water from Day 0 to Day 3. On Days 0, 3, 6, and 9, mice received intraperitoneal injections of 200 μg/mouse αCTLA-4 (BE0131, BioXcell; BE0164, BioXcell) or isotype control IgG (BE0087, BioXcell). Fecal samples were collected daily from mice at the start of DSS drinking for DAI scoring, and terminal dissection was performed on Day 11. For DAI scoring, if the degree of bloody stool is visible to the naked eye, it is scored directly based on severity. If not visible to the naked eye (i.e., occult blood), a fecal occult blood test kit (BA2020B, BaSO) is used for detection, and the degree of occult blood is quantified and scored strictly according to the kit’s standards.

### In vivo treatment

In the DSS-based αCTLA-4-induced colitis model, anti-mouse IL-17A neutralizing antibody (200 μg/mouse; S0B1283, Starter) was administered intraperitoneally at D0, D3, D6, and D9. In the DSS-based αCTLA-4-induced colitis model, MK2206 (p.o.; 30 mg/kg/day; HY-108232, MCE), Rapamycin (i.p.; 5 mg/kg/day; HY-10219, MCE), and 2-DG (i.p.; 500 mg/kg/day; HY-13966, MCE) were administered from D0 to D10. In the colitis tumor-bearing model, mice were administered 1 mg/kg or 3 mg/kg meprednisone (HY-B0260, MCE) and 100 mg/kg or 200 mg/kg metformin (HY-17471A, MCE) from days 3 to 7.

### Immunophenotype analysis

After obtaining the mouse colon, the fat tissue and feces were removed, and the colon was cut lengthwise into small segments. Colon tissues were placed in mucosal layer digestion buffer (10 mL Hank’s + 31 μL 5% (w/v) dithiothreitol (DTT) + 20 μL 0.5 M EDTA + 20 μL FBS) and shaken at 37°C and 220 rpm for 15 min. Then, colon tissues were washed twice with cold PBS. Next, the tissues were minced and placed in 5 mL of the established layer digestion solution (5 mL RPMI 1640 + 2.5 mg trypsin + 7.5 mg collagenase II + 60 μL FBS). The mixture was then shaken at 37°C and 220 rpm for 40 minutes. Subsequently, the digestive fluid was filtered through a 70 μm membrane, then centrifuged, washed, and resuspended to obtain a single-cell suspension. Then the cells were blocked with 4% FBS and anti-CD16/CD32 (BD Biosciences), incubated with surface marker antibodies for 20 minutes at 4℃ and then permeabilized with BD Cytofix/Cytoperm buffer before intracellular labeling antibodies were added for 30 minutes at 4℃. Then transcription factors were labeled for 45 minutes at 4℃ by antibodies after permeabilized with BD TF Fix/Perm buffer. Flow cytometry analysis was performed using ACEA NovoCyte and data processing was done through NovoExpress software. Antibody staining was performed following the manufacturer’s recommendations.

### RNA-Seq

RNA was isolated from fresh colon tissues. Transcriptome libraries were constructed and sequenced by Majorbio. Differential expression was evaluated with DESeq. A fold-change of 2:1 or greater and a false discovery rate (FDR)-corrected p-value of 0.05 or less were set as the threshold for differential genes. KEGG analysis was performed via the MajorBio cloud platform.

### Single-cell sequencing data analysis

Single cell RNA-seq data used in this study were all from the GEO database (GSE144469). All data analysis steps were performed according to the original literature.

### Statistical analysis

The in vivo experiments were randomized but the researchers were not blinded to allocation during experiments and results analysis. Statistical analysis was performed using GraphPad Prism 8 Software. A Student’s t test was used for comparison between the two groups. Multiple comparisons were performed using one-way ANOVA followed by Tukey’s multiple comparisons test or two-way ANOVA followed by Tukey’s multiple comparisons test. Detailed statistical methods and sample sizes in the experiments are described in each figure legend. All statistical tests were two-sided and P-values < 0.05 were considered to be significant. ns not significant; *p < 0.05; **p < 0.01; ***p < 0.001.

## Data availability

All data are available in the main text or the supplementary materials. Correspondence and requests for materials should be addressed to Y.L.

## Acknowledgments

This work was supported by the Shanghai Rising-Star/Sailing Program (24YF2756000). All the schematics are were created with BioRender.com.

## Author contributions

Conceptualization, Y.L. and L.G.; Methodology, Y.L., Z.L., S.W. and R.L.; Formal Analysis, Y.L.; Investigation, Y.L., Z.L., S.W., R.L., R.C, F.L., R.Z., Y.W., C.C., X.Z., F.Q., L.C., Y.Z., and F.D.; Resources, L.G. and Y.L.; Writing – Original Draft, Y.L.; Writing – Review & Editing, L.G. and Y.L.; Supervision, L.G.; Funding Acquisition, L.G. and Y.L.

