## Supplementary Materials for "CTLA-4 inhibitors drive colitis through metabolic reprogramming-mediated Treg/Th17 imbalance"

**This file contains:**

Fig. S1. Direct administration of αCTLA-4 did not induce colitis in mice.

Fig. S2. Establishment of a DSS-based αCTLA-4-mediated colitis.

Fig. S3. Establishment of mouse models of αCLTA-4/αPD-1 combination-mediated colitis and PD-1 inhibitor-mediated colitis.

Fig. S4. Establishment of immuno-humanized mouse models of ICI-mediated colitis.

Fig. S5. The effect of antibody-mediated Treg depletion on αCTLA-4-mediated colitis.

Fig. S6. Evaluation of the effect of metformin on other ICI-mediated colitis.

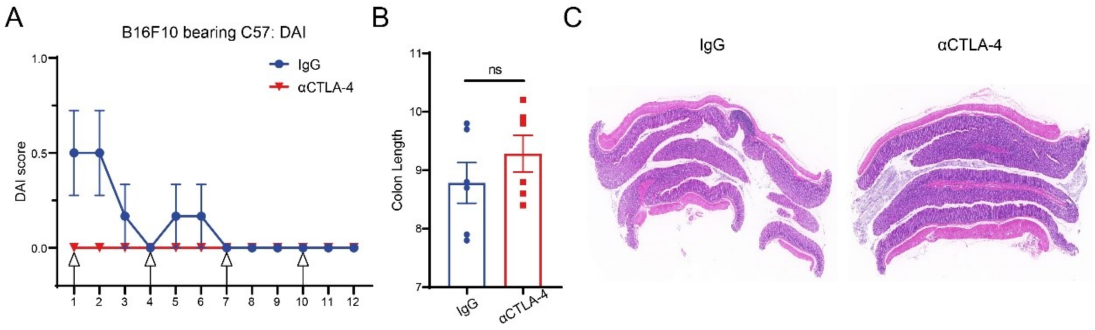


**Fig. S1. Direct administration of αCTLA-4 did not induce colitis in mice.** B16F10 tumor-bearing C57BL/6 mice received four injections of αCTLA-4 9H10 or IgG isotype control. (**A**) Monitoring of DAI scores during the experiment (n=6). (**B**) Colon length of mice at the end of the experiment (n=6). (**C**) Colon length of mice at the end of the experiment (n=6). The data are presented as the mean ± SEM. ns not significant by unpaired t test.

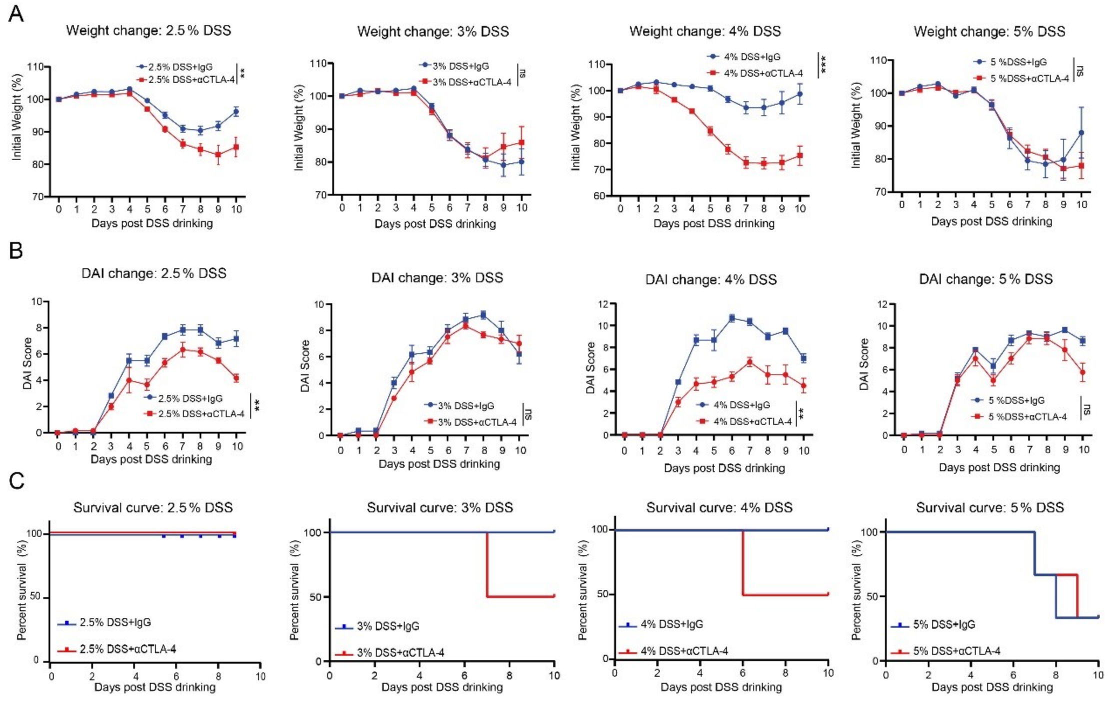


**Fig. S2. Establishment of a DSS-based αCTLA-4-mediated colitis.** B16F10 tumor-bearing C57BL/6 mice were provided with DSS in drinking water at varying concentrations from Day 0 to Day 3. On Days 0, 3, 6, and 9, mice received intraperitoneal injections of 200 μg/mouse αCTLA-4 9H10 or isotype control IgG. (**A**) Weight Monitoring (n=6). (**B**) DAI Score Monitoring (n=6). (**C**) Mouse Survival Monitoring (n=6). The data are presented as the mean ± SEM. ** $p < 0.01$; *** $p < 0.001$; ns not significant by ns not significant by unpaired t test for endpoint.

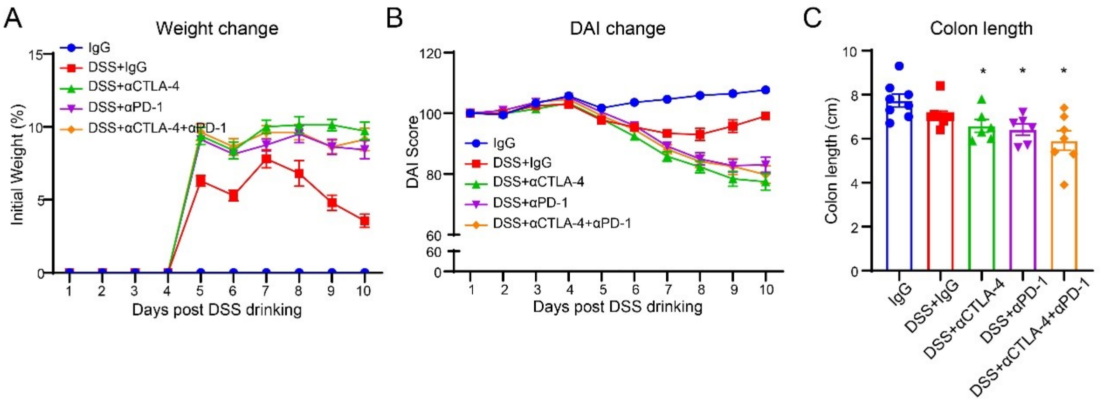


**Fig. S3. Establishment of mouse models of αCLTA-4/αPD-1 combination-mediated colitis and PD-1 inhibitor-mediated colitis.** (**A**) Body weight monitoring (n=6-8). (**B**) DAI score monitoring (n=6-8). (**C**) Mouse colon length (n=6-8). The data are presented as the mean ± SEM. * $p < 0.05$ by one-way ANOVA followed by Tukey's multiple comparisons test.

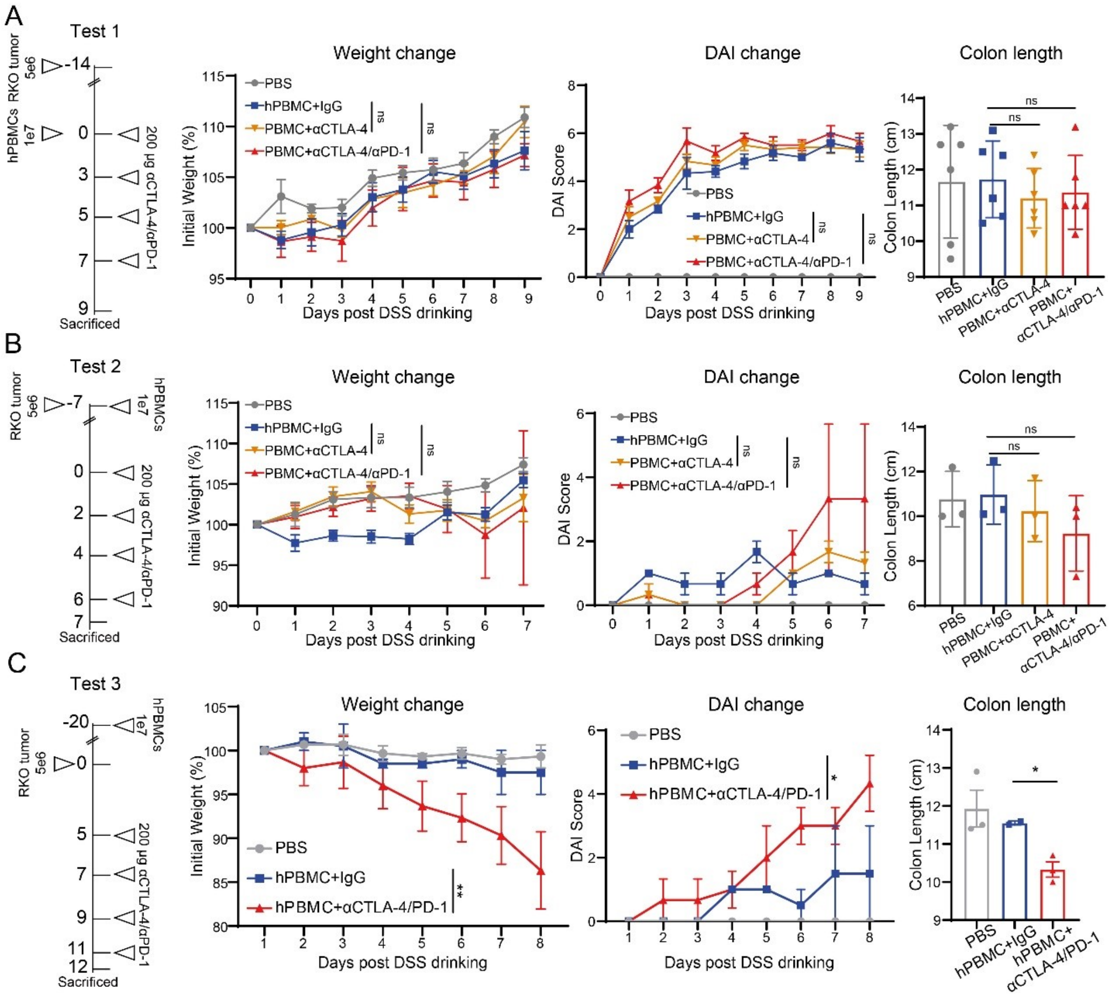


**Fig. S4. Establishment of immuno-humanized mouse models of ICI-mediated colitis.** (**A-C**) Technical roadmaps, body weight monitoring results, DAI monitoring results, and colon length statistics for three different protocols (n=3-6). The data are presented as the mean ± SEM. * $p < 0.05$; ** $p < 0.01$; ns not significant by one-way ANOVA followed by Tukey’s multiple comparisons test.

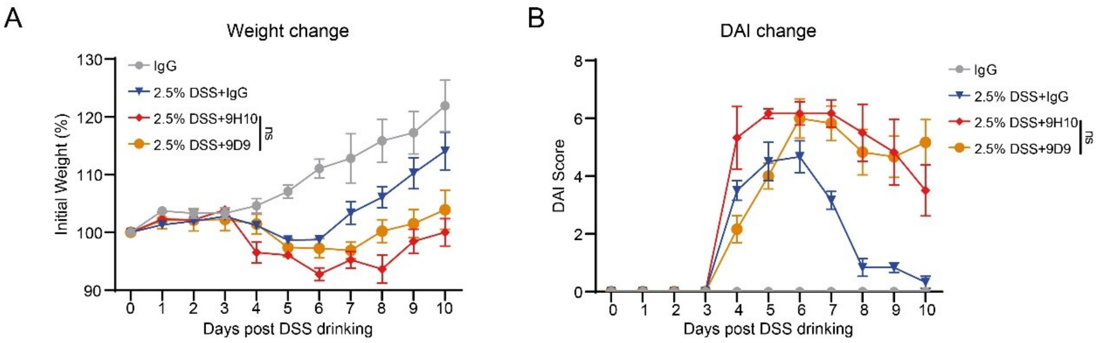


**Fig. S5. The effect of antibody-mediated Treg depletion on αCTLA-4-mediated colitis.** (**A-B**) Body weight monitoring and DAI score monitoring in mice after modeling with two CTLA-4 inhibitors, 9H10 and 9D9 (n=6). The data are presented as the mean ± SEM. ns not significant by one-way ANOVA followed by Tukey's multiple comparisons test.

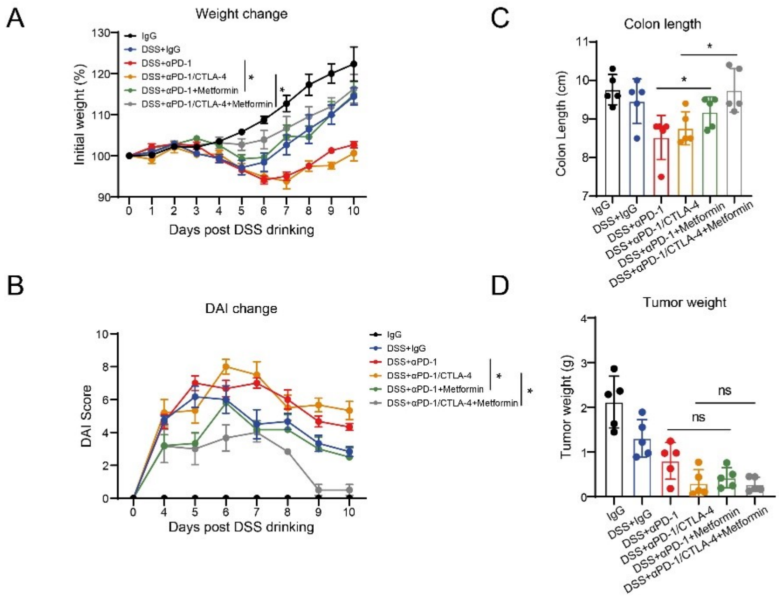


**Fig. S6. Evaluation of the effect of metformin on other ICI-mediated colitis.** (**A-B**) Weight monitoring and DAI score monitoring (n=5). (**C**) Colon Length Measurement (n=5). (**D**) Endpoint tumor weight measurement (n=5). The data are presented as the mean ± SEM. * $p < 0.05$; ns not significant by one-way ANOVA followed by Tukey's multiple comparisons test.
